# NRAMP2 controls manganese partitioning between seed coat and embryo to regulate seed vigour

**DOI:** 10.64898/2026.03.04.709681

**Authors:** Alexandra Leskova, Marie-Pierre Isaure, Tou Cheu Xiong, Camille Rivard, Hiram Castillo-Michel, Kadda Mejoubi, Andrea Somogyi, Sandrine Chay, Katarzyna Bierla, Stéphane Mari, Catherine Curie

## Abstract

Seed development and germination depend on the coordinated transport of macronutrients and micronutrients, which sustain the embryo and determine seed vigour. Manganese (Mn) is an essential micronutrient involved in various metabolic processes, yet its role in seed development is not well understood. In this study, we investigated manganese (Mn) homeostasis in seeds and identified NRAMP2, a previously characterised trans-Golgi-localised Mn exporter, as a key transporter that controls Mn delivery to the embryo. Promoter analysis showed NRAMP2 expression in the chalazal region, which is a central hub for the exchange of nutrients between maternal tissues and the developing seed. X-ray fluorescence mapping revealed mislocalisation of Mn in *nramp2* mutant seeds, retention of Mn on the seed coat and markedly reduced levels of Mn in the embryo. Functionally, Mn depletion delayed seed germination and increased physiological dormancy by reducing reactive oxygen species (ROS) accumulation. Our results uncover the vital role of Mn in supporting embryo development and seed vigour, establishing NRAMP2 as a pivotal component of the Mn transport machinery during seed development.

## INTRODUCTION

Seeds serve as an important food source for humans and animals. Seed development requires an optimal flow of assimilates and a cocktail of macro- and micronutrients to feed the embryo. Enrichment by metallic micronutrients in particular, such as manganese (Mn), iron (Fe) or zinc (Zn) is unequivocal to combat nutrient deficiency, aka "hidden hunger," that has been linked to various malnutrition-related human diseases, including anemia and neurological disorders (reviewed in (*1*)). Nutrient transport towards the seeds begins in the plant’s conducting tissues and proceeds to the funiculus, which connects maternal tissues to the seed. At the terminal end of the phloem, the unloading domain, so-called chalazal seed coat, nutrients are redistributed across several cell layers into the embryo via symplastic (transporter-independent) and apoplastic (transporter-dependent) pathways. Symplastic pathways dominate the unloading domain due to the high density of plasmodesmata; however, numerous transporter proteins for amino acids and glucosinolates, are also expressed in this zone (*2*, *3*). Apoplastic transport processes within seeds are relatively well-characterized for sucrose, amino acids, and some specialized metabolites, relying on H⁺-symporters for uptake and facilitators for bidirectional flux driven by concentration gradients (*2*).

In contrast, transporter-driven uptake of essential metals towards the embryo tissues remain poorly documented. Notable examples include the P1B type ATPases HMA2 and HMA4, which facilitate Zn transfer from the seed coat to the embryo in Arabidopsis (*4*). In monocotyledonous plants, ZmYSL2 and OsYSL9 are responsible for the transport of Fe complexed with nicotianamine from the endosperm towards the embryo (*5*, *6*). At the intracellular level, tonoplast-localized transporters such as OsVIT2 and TaVIT2 in aleurone cells and AtVIT1 in embryonic cell files promote Fe sequestration in vacuolar globoids (*7–9*). Manganese transport within embryos is mediated by the tonoplastic transporter MTP8, which can even substitute for Fe transport in the absence of VIT1, while conversely VIT1 is responsible for Mn loading into vacuoles when MTP8 is missing (*9*, *10*).

Nutrient deficiencies impact seed viability to varying degrees. Phosphate limitation exerts relatively minor effects on germination, whereas maternal nitrate availability correlates positively with seed vigor (*11*). Likewise, severe Fe limitation prolongs seed dormancy across multiple Arabidopsis accession lines (*12*). Despite its essential roles, the influence of Mn on seed physiology and agricultural traits remains much less explored. Evidence suggests that MTP8-mediated vacuolar sequestration of Mn in cortical and subepidermal layers of the embryo during development and germination is critical; *mtp8* seeds from low-Mn fed plants or imbibed with excess Mn show increased seed dormancy (*10*). Yet, how Mn regulate seed development or viability remains unknown.

Mn serves as a cofactor for numerous enzymes and thus potentially governs diverse processes contributing to seed viability. It supports reactive oxygen species (ROS) metabolism through Mn-dependent superoxide dismutases, which convert superoxide to hydrogen peroxide in developing and germinating seeds. Mn is furthermore a cofactor of multiple enzymes involved in synthesizing cell wall components—primarily pectins, the loosening of which in the endosperm and embryo are essential for radicle protrusion and emergence during germination.

To investigate Mn import into plant embryos, we examined NRAMP2, a seed-expressed Mn transporter and the impact of Mn misallocation on seed development and vigor. NRAMP transporters (SLC11 family), first identified for their role in animals innate immunity where they mediate proton-coupled Fe^2+/^Mn^2+^ efflux from phagosomes to starve pathogens (reviewed in (*13*)), are ancient metal-handling systems conserved across kingdoms. In planta NRAMP transporters mediate Fe, Mn and Cd transport, among them several are implicated in seed metal transport. The plasma membrane-localized NRAMP1 is expressed at the seed coat and potentially mediates Fe delivery to the embryo (*14*). Loss of its negative regulator INNER NO OUTER (INO) triggers excessive ROS accumulation due to NRAMP1-driven Fe overaccumulation. Tonoplast-localized NRAMPs, such as rice NRAMP2 and the Arabidopsis NRAMP3 and NRAMP4, facilitate Fe efflux from vacuoles during germination (*15*, *16*).We and others have previously demonstrated that NRAMP2 pumps Mn out of the trans-Golgi network to feed various organelles with Mn (*17*, *18*). In this study we show that NRAMP2 functions as a Mn transporter in the chalazal region of the seed, from where it distributes Mn to the embryo. X-ray fluorescence analyses demonstrate mislocalization of Mn in the absence of NRAMP2. Moreover, reduced levels of Mn in the embryo enhance physiological dormancy due to decreased ROS accumulation, thereby uncovering a critical role for Mn in processes essential to seed vigor.

## RESULTS AND DISCUSSION

### The chalazal seed coat-localised NRAMP2 is responsible for Mn delivery into the embryo

We and others have previously shown that *NRAMP2* is expressed in the vegetative tissues throughout plant development (*17*, *18*). Publicly available expression databases suggested that *NRAMP2* is transcribed in seed tissues as well, particularly in the chalazal region and to less extent in general seed coat (*19–21*)**; Supplementary Fig. 1**). Leveraging our transcriptional fusion lines of *NRAMP2* described in Alejandro et al. (*17*), we verified the tissue-specific expression patterns in developing Arabidopsis seeds. We observed GUS activity in two hotspots: in the funiculus, which attaches the developing seed to the maternal tissues, and in the chalazal seed coat domain within the seeds (**Fig. 1**). Of note is the junction of the phloem pole within the GUS-expressing chalazal domain (**Fig. 1f**). We cannot exclude however the possibility that the expression territory of *NRAMP2* extends to the general seed coat.

**Figure 1.**
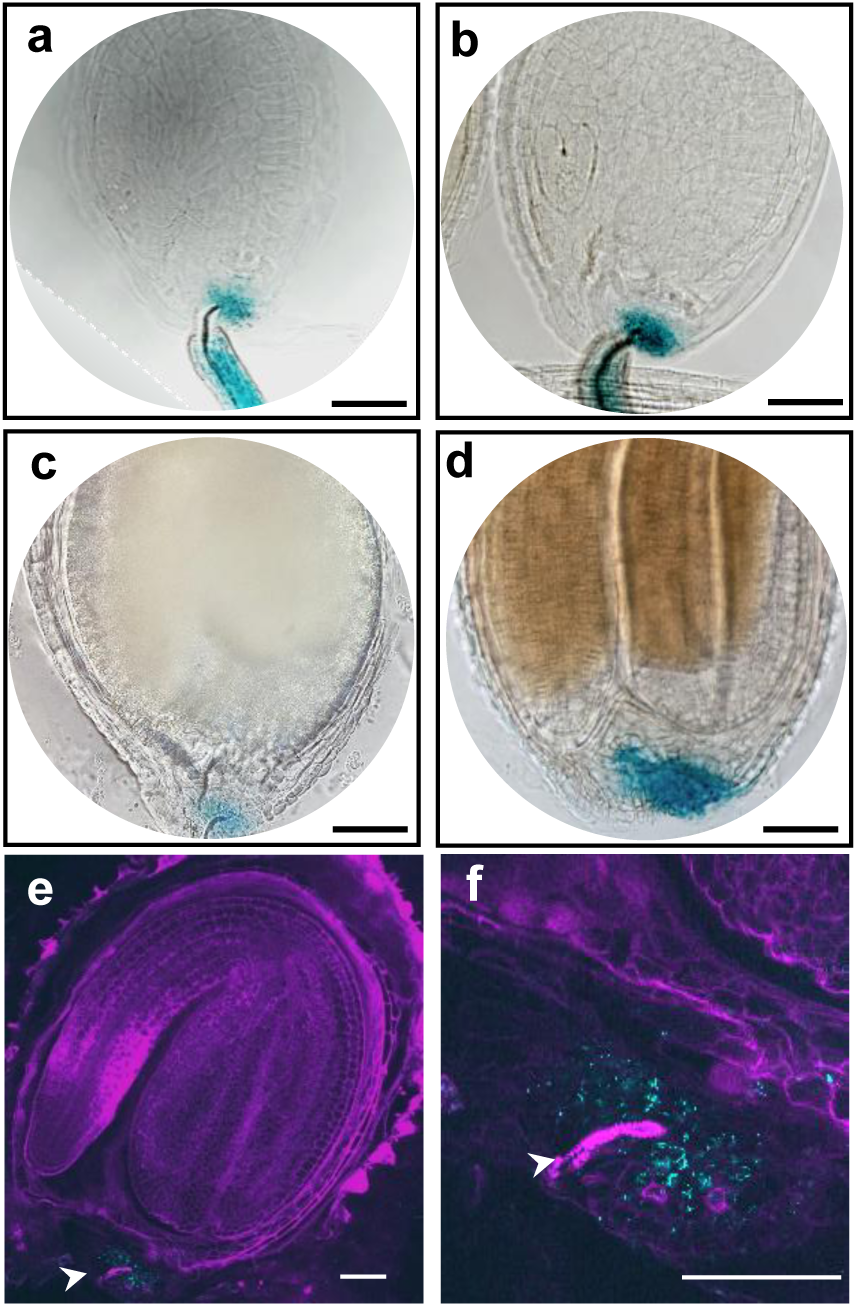
**NRAMP2 is expressed in the funiculus and chalazal seed coat of the seeds**. *pNRAMP2::GUS* activity in seeds of heart (**a**), torpedo (**b**), late bent cotyledon (**c**) and mature green (**d**) stage. **e**, GUS reflection image of mature seeds, **f,** inset of chalazal seed coat region, cyan – GUS reflection, magenta – propidium iodide counterstain. Arrowhead points to the phloem end pole. Scalebars = 50 µm.

To investigate the function of NRAMP2, we previously studied two knock-down mutants of *nramp2* (*17*). In this study, to explore the function of NRAMP2 in seeds, we generated two additional *nramp2* mutant alleles in Col-0 background by CRISPR-Cas9 (**Supplementary Fig. 2**). Both the *nramp2-5* and *nramp2-6* alleles exhibited retarded root growth and decrease in shoot biomass specifically under Mn deficiency (**Supplementary Fig. 3a and b)**, phenocopying the previously described EMS mutant *nramp2-1* (*18*).

Given that NRAMP2 functions as a Mn transporter, we wondered whether lack of *NRAMP2* expression could alter Mn accumulation and/or distribution in the seeds. Elemental profiling of whole dry seeds revealed a significant 30% decrease of Mn content and a minor, 10% decrease of zinc (Zn) content in *nramp2* and no change in Fe, compared to wild-type seeds (**Supplementary Fig. 4a-c**). No significant change in the amount of other metals or nutrients was observed between wild-type and mutant seeds (**Supplementary Table I**). Given that NRAMP2 is expressed in the vegetative parts of the plants, such as roots and shoots (*17*, *18*) as well as funiculus (this study), changes in total Mn content in the seeds could be the final consequence of disturbed Mn transport throughout plant growth. Supplying high doses of Mn during plant development increased the total Mn content of seeds regardless of the genotype, but, more importantly, it erased the difference in Mn content between *nramp2* and wild-type seeds. This is indicative of the role of NRAMP2 in transporting Mn to seeds (see **Supplementary Fig. 4D**).

Next, to investigate whether NRAMP2-mediated Mn transport contributes to Mn distribution among maternal and embryonic seed tissues, we performed spatial mapping of metals in seeds. Firstly, we employed X-ray fluorescence tomography of intact dry seeds. Single slice and 3D sparse tomography reconstruction of whole seeds showed a dramatic difference in Mn distribution between wild-type and *nramp2-6* **(Fig. 2a-f).** As previously reported, the majority of Mn in wild-type plants was confined to the embryo, specifically the cortical layer of the hypocotyls and the sub-epidermal layer on the abaxial side of the cotyledons (*9*, *10*) (**Fig. 2b,c**). In contrast, only a small amount of Mn could be detected in the embryos of *nramp2-6*, with Mn accumulating almost exclusively at the outer cell layers of the seeds (**Fig. 2e,f**). By comparison, we found no difference in Fe distribution between wild-type and the mutant (**Fig. 2a-f**).

**Figure 2.**
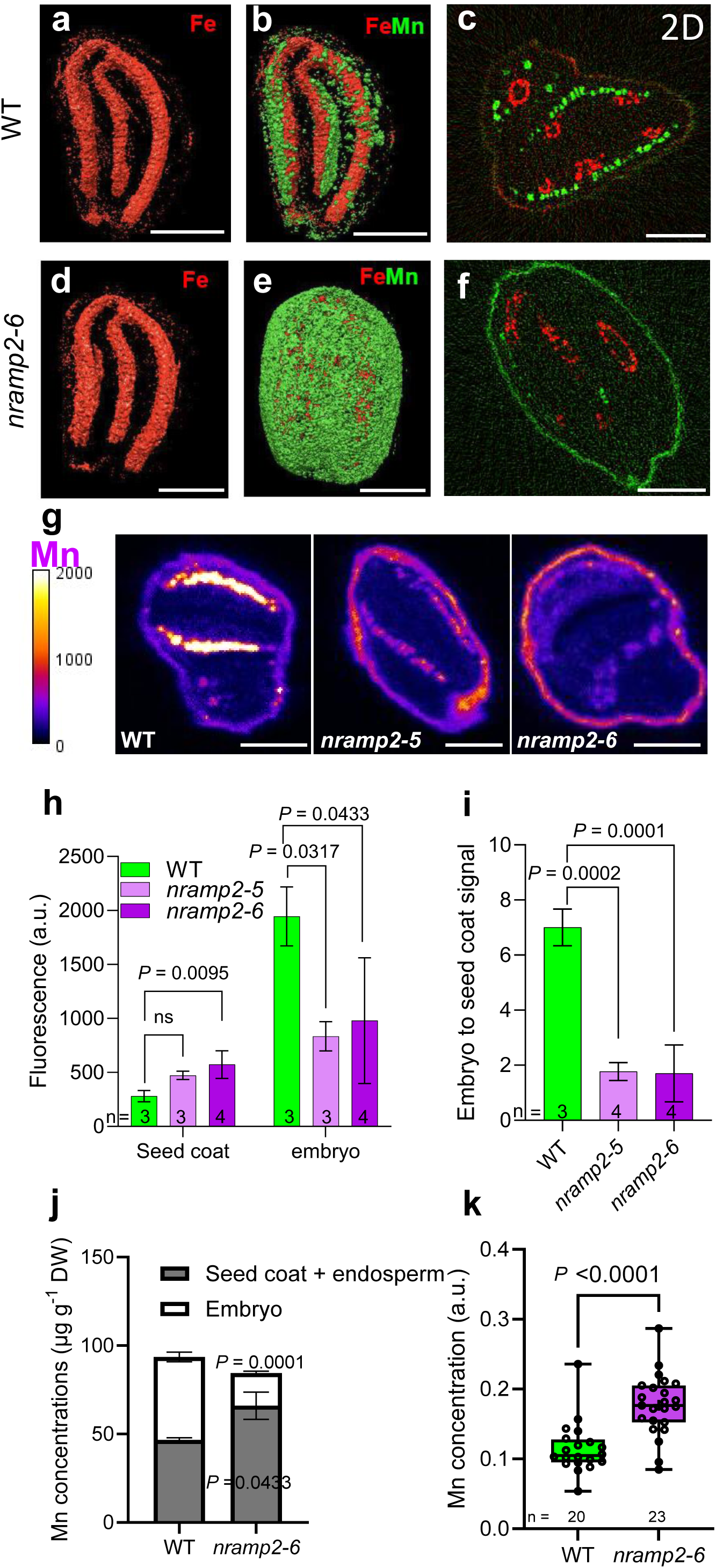
NRAMP2 regulates Mn loading from seed coat to embryo. Spatial mapping of manganese in dry seeds using micro-X-ray fluorescence imaging. **a-b, d-e**, 3D sparse tomogaphy reconstruction of Fe (**a,d**), Mn overlayed with Fe (**b,e**). **c,f**, reconstructed high resolution 2D single slice tomograms of Fe and Mn using NANOSCOPIUM beamline at Synchrotron Soleil, France. **a-c,** wild type, **d-f,** *nramp2-6* seeds. **g,** Micro-X-ray fluorescence imaging of Mn in dry seed sections using Lucia beamline at Synchrotron Soleil. Representative images of seed sections are shown. **h,i,** Quantification of fluorescence signal in regions of interests within the embryo and the outer cell layer. Bars represent means ± SD of three independent seed sections. *P*-values were determined by one-way ANNOVA and followed by post-hoc Tukey’s test. **j,** Mn contents in one-hour imbibed seeds dissected into embryos and seed coat+endosperm, measured by ICP-OES. Bars represent means ± SD of three independent samples bulked from around 400 seeds. *P*-values indicated in the bars were determined by two-tailed unpaired Student’s *t* test between wild type and *nramp2-6* for each seed section. **k,** Accumulation of Mn in laser-ablated envelopes of dry seeds, quantified by ICP-MS. Box plot represents all data points, whiskers the 25th percentile, median and 75th percentile. The *P*-value was determined by two-tailed unpaired Student’s *t* test. Scalebars = 100 µm.

To strengthen these results and perform a semi-quantitative analysis of metal levels in seed tissues, we employed a 2D micro-X-ray-fluorescence imaging of Mn and other elements in dry seed sections of the wild-type and the two alleles of *nramp2* **(Fig. 2g**). We quantified the fluorescence signal intensities from regions of interest of the embryo and outer cell layers of multiple seed sections. Compared to the wild-type, Mn abundance in the outer cell layer increased by approximately 32% in *nramp2-5* and 47% in *nramp2-6*, with statistical significance observed only for *nramp2-6* (**Fig. 2h**). Conversely, the embryos of the *nramp2-5* and *nramp2-6* mutant alleles were severely Mn-depleted, by 53% and 47%, respectively, compared to the wild type (**Fig. 2h**), resulting in dramatic reduction of the embryonic-to-seed coat Mn ratio in the mutants (**Fig. 2i**). In contrast to Mn, the mutation did not affect the partitioning of any of the other abundant macroelements, such as calcium, sulphur or phosphorus (**Supplementary Fig. 5**).

To confirm the misdistribution of Mn between the envelope and embryo in the *nramp2* mutant, we undertook two complementary approaches. First, we dissected imbibed seeds into embryos and seed coat/endosperm and analysed Mn content in each fraction. Consistent with ray mapping results, Mn levels were about 30% higher in the envelope of *nramp2-6*, whereas the embryo fraction contained around 60% compared with wild-type (**Fig. 2j**).

Second, we performed laser ablation of the seed coat, coupled to Inductively coupled plasma mass spectrometry (ICP-MS), to determine whether it is indeed the seed coat that overaccumulates Mn in the *nramp2* mutant. Consistent with results of X-ray fluorescence imaging and seed dissection analyses, the ablated seed coat of *nramp2-6* accumulated 35% more Mn than that of the wild-type (**Fig. 2k**). Taken together, these results demonstrate that NRAMP2 controls the uptake of Mn into the seeds and its transport from the seed coat towards the embryo.

### NRAMP2 mediates Mn delivery to subepidermal—but not endodermal—layers of mature embryos

To gain further insight into the timing and mechanism of NRAMP2-mediated Mn transport in seeds, we investigated metal distribution patterns during seed development, in the earlier bent cotyledon stage and the later mature seed stage using synchrotron X-ray fluorescence imaging. Technical limitations prevented imaging of intact seeds prior to the bent cotyledon stage and reliable preservation of seed coats (including the chalazal region) at all stages. Embryos, however, were well preserved from the bent cotyledon stage onward following plunge freezing in liquid nitrogen. In wild-type embryos, Zn and Fe accumulation patterns remained consistent between bent cotyledon and mature green stages, with Zn localising across all embryo layers and Fe accumulating in the endodermis (**Fig. 3a,b,e,f**)—mirroring patterns in dry seeds (**Fig. 2a**; (*10*)).

**Figure 3.**
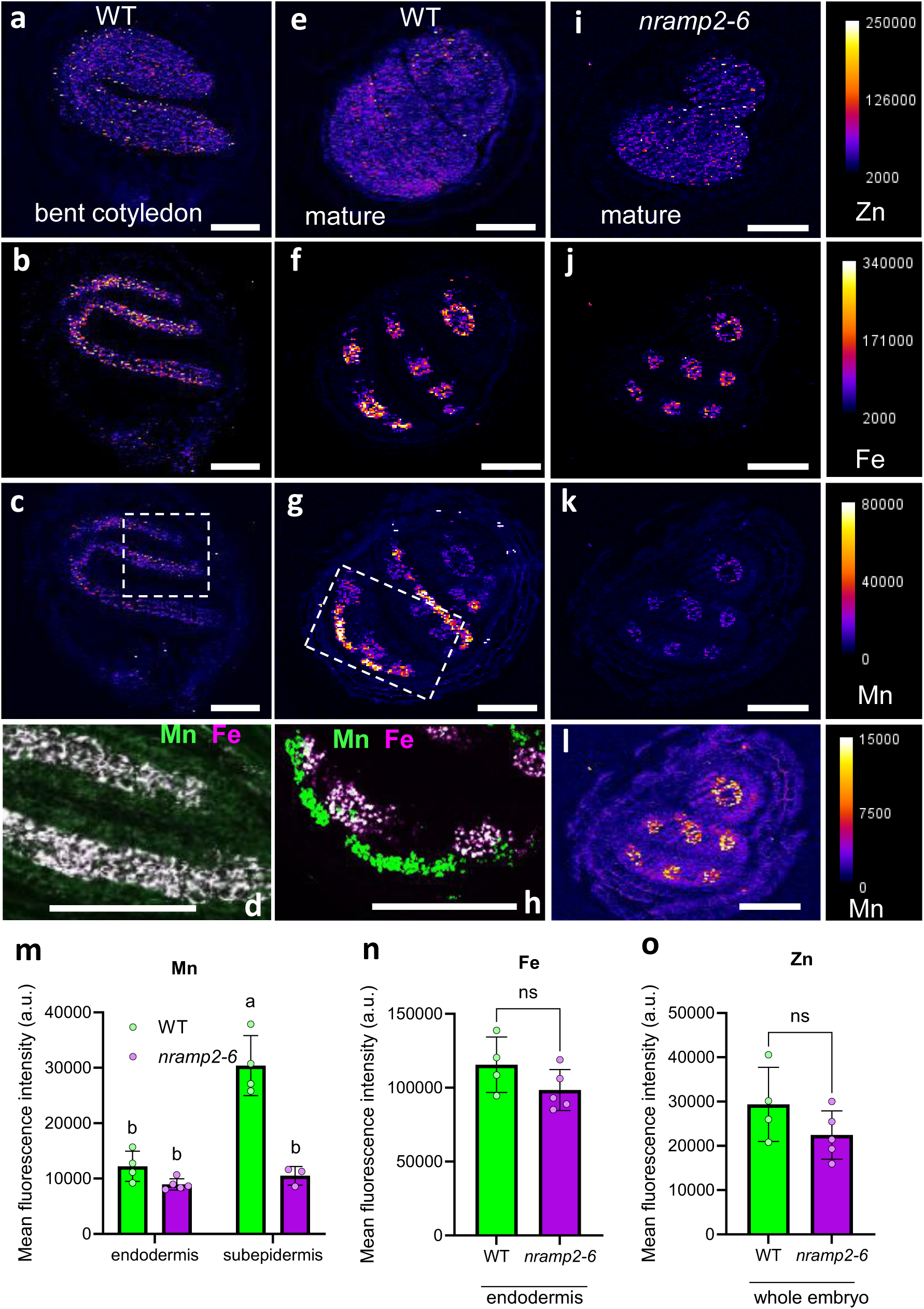
Manganese accumulation in embryonic subepidermal layers is NRAMP2-dependent. Synchrotron X-ray fluorescence mapping of Zn **(a,e,i**), Fe (**b,f,j**) and Mn (**c,g,k,l**) in wild type bent cotyledon and wild type and *nramp2-6* mature green stages using ID21 beamline of Synchrotron ESRF, France. **a-d**, wild type, bent cotyledon stage, **e-h**, Wild type mature green stage, **i-l**, *nramp2-6* mature green stage. **d,e,** High resolution overlay of Mn (green) and Fe (magenta) in bent cotyledon and mature cotyledons, outlined in **c** and **g** images, respectively. **m,n,o**, Quantification of mean pixel intensities of Mn, Fe and Zn X-ray fluorescence in embryonic layers of mature seeds. Bars represent means ± SD of sections of 4-5 independent seeds. Different letters indicate significant differences (one-way ANOVA followed by post-hoc Tukey’s test, *P* < 0.05). Scalebars = 100 µm.

Unexpectedly, however, at the bent-cotyledon stage, Mn exhibited negligible accumulation in its ultimate target tissues, i.e. the subepidermal layers of cotyledons and hypocotyl cortex. Instead, Mn accumulated predominantly in the endodermal layer of the embryo, colocalizing with Fe (**Fig. 3c,d**). By contrast, at the mature green stage, large amounts of Mn were detected in subepidermal layers, matching the localization observed in dry seed embryos **Fig. 3g,h and Supplementary Fig. 6a,b**). The quantitatively less abundant, endodermal pool of Mn persisted to comparable amounts in the mature embryos, indicating that subepidermal Mn in mature seeds represents a distinct, newly acquired Mn pool rather than being remobilized from endodermal stores. The late establishment of the major pool of Mn is consistent with an earlier report showing that Mn transfer from the seed coat/endosperm to the embryo proceeds more slowly than that of other metals such as Zn (*22*).

Notably, mutating NRAMP2 selectively depleted the subepidermal Mn pool of mature embryos, while Mn levels in the endodermis and the overall distribution of other metals remained unchanged **(Fig. 3i-o, Supplementary Fig. 7**). Together these observations strongly suggest the existence of two sequential import pathways into Arabidopsis embryo, operating at different developmental stages and likely involving distinct sets of transporters. Early in development, Mn provision to the embryo does not depend on chalaza-localized NRAMP2, implying either the involvement of another chalazal transporter or the use of an alternative transport route. Once reaching the embryos, the vacuolar Fe transporter VIT1 represents a strong candidate for mediating Mn sequestration in endodermal vacuoles of the early-stage embryos, as it is capable of Mn transport in yeast (*9*), and is expressed as early as the torpedo stage (*10*).

Later in seed development, *nramp2* mutants fail to accumulate Mn in subepidermal embryo tissues, indicating that NRAMP2 governs the formation of the large Mn pool that builds up in this region. The Mn transporter MTP8, previously shown to sequester Mn into the vacuoles of subepidermal cotyledon and cortical hypocotyl cells, is expressed later during seed development, similar to NRAMP2 ((*10*); **Supplementary Fig. 1**). MTP8 therefore represents a good candidate to act in concert with NRAMP2 to promote Mn accumulation in the vacuoles of cortical and subepidermal cells in mature embryos.

The ectopic pool of Mn in the seed coat of *nramp2* suggests that Mn reaching the chalazal seed coat in the mutant diffuses laterally into the first two outer seed coat layers, which are symplastically connected. This abnormal Mn retention provides compelling evidence that active Mn transport by NRAMP2 is required and/or is a prerequisite of Mn transfer within symplastically separated layers of seed coat and between seed coat and endosperm. Interestingly, similar to amino acid transporters, seed coat-expressed Fe/Mn transporters - such as NRAMP1 - operate even in symplastically connected outer integuments of the seed coat (*14*, *23*). Thus, transporters in conjunction with symplastic ion diffusion pathways mediate the intricate transport of metals in seeds.

### Ectopic accumulation of Mn in the seed coat of *nramp2* does not affect its anatomical or morphological properties

Next, we sought to assess the impact of insufficient Mn delivery into the embryos on seed development and postembryonic growth. Both developing and dry seeds of the *nramp2* mutant harboured normal size and weight, indistinguishable from wild type (**Supplementary Fig. 8a-e**). Mn serves as an essential cofactor of Golgi-localized glycosyl transferases, and is crucial for the biosynthesis of multiple cell wall components, such as pectin and, to a lesser extent cellulose (reviewed in (*24*, *25*)). We thus hypothesized that ectopic overaccumulation of Mn in the seed coat of *nramp2* would affect its properties. To test this, we examined the seed coat ultrastructure and permeability, as well as pectin and cellulose deposition. However, none of these parameters differed between wild-type and *nramp2* seeds (**Supplementary Fig. 9a-d).** We had previously shown that the cutin layer diminishes in Arabidopsis leaves of the *nramp2* mutant under Mn deficiency (*17*). Thus, as a readout of cuticle integrity, we tested the permeability of isolated and fully expanded embryos (11 DAP) to toluidine blue (*26*). Unlike the cutin synthesis *bdg-2* mutant, which showed increased dye uptake, *nramp2-6* embryos were indistinguishable from the wild-type (**Supplementary Fig. 9e**).

### NRAMP2-mediated Mn transport to the embryo positively regulates accumulation of reactive oxygen species to release dormancy

Proper Mn distribution within the seed embryo is crucial for germination efficiency (*10*). We therefore examined whether the sparse embryonic Mn reserves in *nramp2* could impact seed vigour. Although NRAMP2 is not expressed in imbibed or germinated seeds, we hypothesized that misallocation of Mn in seeds could nonetheless compromise subsequent germination performance. We thus tested seed germination efficiency of *nramp2* **(Fig. 4a).** The proportion of germinating *nramp2-6* seeds decreased by approximately 50% compared to the wild type. Supplementation of Mn to the germinating medium restored the germination efficiency of *nramp2-6* seeds to wild-type levels (**Fig. 4a).** Because inactivation of NRAMP2 did not measurably affect seed coat structure, a seed coat–imposed barrier is unlikely to explain the reduced germination of *nramp2* seeds. We therefore hypothesized that reduced Mn accumulation in the embryo could influence physiological dormancy. Prolonged cold incubation of *nramp2* seeds rescued the germination defect of *nramp2* to wild-type levels (**Fig. 4b).** The finding that *nramp2* seeds exhibit enhanced physiological dormancy supports a role for NRAMP2-dependent Mn allocation in dormancy regulation.

**Figure 4.**
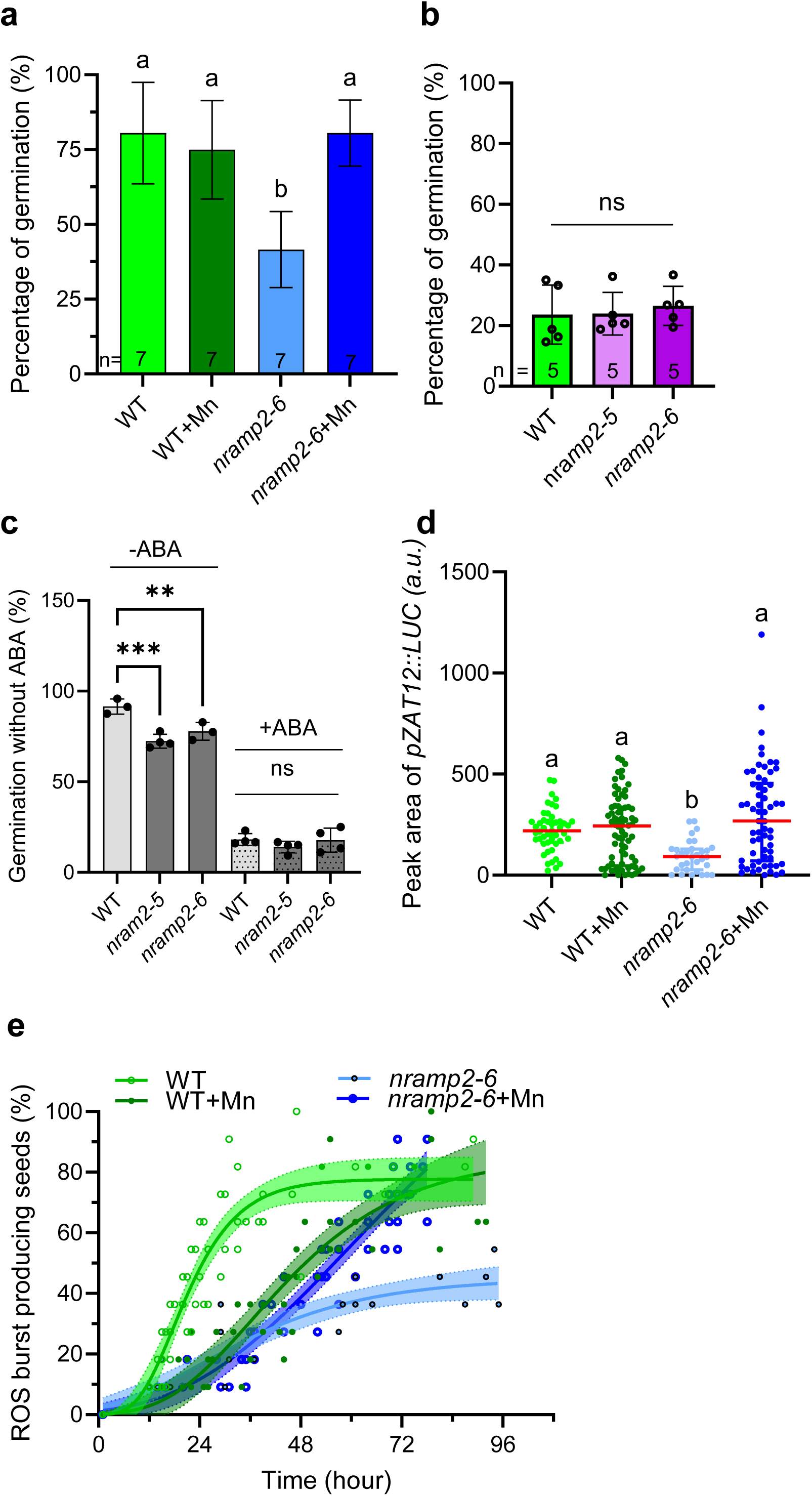
Inefficient Mn transport by NRAMP2 renders seeds more dormant. **a**, Percentage of seed germination in dark without cold stratification. Bars show means ± SD of 7 biological replicates consisting of 10 seeds/replicate. *P*-values were determined by one-way ANNOVA and followed by post-hoc Tukey’s test. **b,** Seed germination in dark and 4°C for 14 days. Bars show means ± SD of 5 biological replicates. Statistical significance was determined by one-way ANNOVA and followed by post-hoc Tukey’s test. **c,** Seed germination with or without added (2 µM) abscisic acid (ABA). **d**, Peak area of *pZAT12::LUC* luminescence calculated during 96 hours after sowing. The scatter dot plot displays the median with interquartile range consisting of 32-62 biological replicates. Different letters indicate significant differences (non-parametric test, followed by post-hoc Kruskal-Wallis test, *P* < 0.05). **e**, Percentage of *pZAT12::LUC* seeds producing ROS burst. The curves are fitted according to Gompertz function. Seeds were sown on 0.5% agar buffered at pH 6 with MES with luciferin 250µM supplemented or not with 500 µM MnSO_4_ and germinated for 96 hours.

Since abscisic acid (ABA) is a central regulator of seed dormancy, we next tested whether the seeds of the *nramp2* mutant display altered sensitivity to exogenous ABA. Application of ABA (2µM) suppressed germination in both wild-type and *nramp2* seeds, with no significant difference between genotypes (**Fig. 4c**), thus ruling out a role of the ABA pathway in the increased dormancy observed in *nramp2* seeds. Rather than reflecting a defect in ABA signalling *per se*, the dormancy phenotype of *nramp2* may arise from ABA-independent mechanisms or from altered hormonal balance upstream or parallel to ABA action.

Since reactive oxygen species (ROS) production precedes germination and Mn modulates ROS homeostasis, we wondered if *nramp2* seeds has inherently altered ROS contents. To this end, we introduced the luminescent transcriptional ROS signalling marker *pZAT12::LUC* into the *nramp2-6* background and monitored the luminescence signal *in vivo* during germination in the dark. A sharp increase of luminescence signal, indicative of a ROS burst, was observed exclusively in germinating seeds, regardless of the genotype. In contrast to wild-type, germinating *nramp2-6* seeds manifested significantly reduced ROS production when germinated on 0 Mn over the course of the experiment **(Fig. 4d).** To test whether exogenous Mn could rescue ROS dynamics and the dormancy phenotype, we supplemented the growth medium with Mn (500 µM). Within 4 days, excess Mn significantly enhanced ROS production of germinated *nramp2* seeds (**Fig. 4d**). Moreover, the proportion of ROS burst-producing *nramp2-6* seeds increased with Mn supply in comparison to wild type (**Fig. 4e**), correlating with improved germination efficiency (**Fig. 4a**).

NRAMP2 likely ensures Mn allocation to the embryo during seed development as a prerequisite for timely ROS production and dormancy release. Since the ZAT12 promoter and expression responds to H_2_O_2_ and particularly to superoxide (*27*, *28*), we assumed that superoxide dismutation via Mn-SOD would be affected in the Mn-deficient *nramp2* seeds. We quantified SOD isoforms in mature seed stage when Mn-SOD peaks (*22*). However mature *nramp2-6* seeds showed normal MnSOD activity **(Supplementary Fig. 10),** consistent with reports that Mn deficiency does not alter MnSOD in higher plants (*17*). Thus, the precise component of ROS producing machinery affected in *nramp2* seeds remains to be identified.

Mn serves as a cofactor for several enzymes that have been previously directly or indirectly implicated in seed dormancy regulation. PP2C-type phosphatases, such as PAPP2C require Mg²⁺ or Mn²⁺ for activity and are involved in dephosphorylation of known dormancy regulators, phytochromes and phytochrome-interacting proteins (*29*). Another PP2C member, IBO, when mutated manifests germination defects as well, although exact implication of IBO in dormancy is yet unknown (*30*). Additionally, Mn is a cofactor of arginine hydrolysing enzyme - L-arginine amidinohydrolase (arginases), highly expressed during germination. The mutation of arginases leads to activation of alternate pathways of arginine catabolism, resulting in elevated nitric oxide levels, reduced ROS accumulation, and impacted seed germination (*31–33*). Whether these or other enzymes are misregulated in Mn deficient *nramp2* are to be further investigated.

## CONCLUSIONS

We demonstrate that Mn delivery into and within the embryo is spatially and timely regulated in developing seeds to ensure adequate embryonic Mn accumulation and to support seed viability. Considerable Mn redistribution from the seed coat and endosperm happens at the rather late bent cotyledon stage (*22*), supplying Mn to endodermal vacuoles of the embryo via yet unidentified plasma membrane influx/efflux transporters, as well as vacuolar VIT1 and MTP8 transporters (*9*, *10*). In mature seeds, NRAMP2 aids in the transport of newly arriving Mn from the chalazal and general seed coat into the endosperm. Mn is then further loaded to the embryo by unknown transporters and retained in subepidermal embryonic vacuoles via the MTP8 vacuolar transporter (*10*) to ensure proper ROS production and successful seed germination.

The specific role of NRAMP2 in mediating Mn shuttling at the chalazal seed coat during late developmental stages may be linked to the differentiation status of neighboring tissues: (i) the chalazal endosperm transitions from a syncytial, transfer cell-like structure, to a cellularised state at the bent cotyledon stage, overlapping with (ii) the degradation of proliferative chalazal tissues at the boundary between the chalazal endosperm and seed coat (*34*). These developmental transitions coincide with a massive loading of Mn into the embryo (this study). In mature seeds, where remnants of the maternal nucellus have disappeared, direct symplastic connections between the chalazal seed coat and endosperm are lost. It is therefore plausible that nutrient transfer, between these tissues at maturity increasingly depends on apoplastic transport mechanisms. This is also consistent with the fact that Mn is still able to be transported into the general seed coat in the mutant, probably via symplastic routes.

At the subcellular level, however, how can the role of NRAMP2 in cell-to-cell Mn transport be reconciled with its previously reported function as a trans-Golgi-localized Mn efflux transporter? A comparable case exists for the Golgi/trans-Golgi–localized phosphate exporter PHO1, which mediates phosphate movement from the chalazal seed coat to the embryo during seed development (*35*). Recently, PHO1 stabilisation at the plasma membrane by inhibited clathrin-mediated endocytosis (CME) was shown to reduce its phosphate export activity into the apoplast of xylem parenchyma (*36*). Whether PHO1 actually imports phosphate into post-Golgi structures that are subsequently secreted via exocytosis remains unresolved (*36*). Other Golgi or trans-Golgi network (TGN) transporters, such as MTP11 or nicotianamine-transporting vesicular proteins, may operate in a similar fashion to mediate intercellular substrate movement through vesicular secretion (*37–40*). Evidence of Fe accumulation in vesicles across diverse wheat grain tissues further underscores the potential importance of vesicular metal transport mechanisms (*41*). If NRAMP2-containing vesicles are targeted for exocytosis, NRAMP2 could restrict Mn export from the cell, whereas in the *nramp2* mutant, where more Mn is trapped in these secretory vesicles, more Mn might diffuse into the apoplast. In this scenario, Mn would accumulate within the apoplastic space of the seed coat. High-resolution subcellular imaging approaches such as STEM-EDX or nano X-ray techniques of well-preserved seeds would be necessary to prove the mode of action of TGN-localised NRAMP2 transporter. In conclusion, enhancing embryonic Mn contents via genetic engineering of seed coat transporters represents a powerful strategy to enhance seed vigor.

## Material and methods

### Plant material and seed harvest

Seeds of wild type Col-0 *Arabidopsis thaliana*, *bdg-2* (*42*), *nramp2-5, nramp2-6* CRISPR Cas9 mutants, described in Supplementary Figure 2, *nramp2-1* (*18*), *pNRAMP2::GUS* (*17*) and *pZAT12::LUC* line described in (*27*), was crossed with *nramp2-6* allele and were used. Dry and mature seeds were harvested from plants grown in soil substrates of greenhouse, fertilised regularly and with controlled photoperiod of 16-light/8-h dark cycle.

### Seed germination assays

About 2 weeks to 3 months after-ripened seeds were sown onto 0.5% (w/v) agar plates, with pH 6 adjusted with 2.5 mM MES-KOH, with or without 500 µM MnSO_4_.H_2_O. The seeds were sown under low intensity light (≈20 μmol.m^-2^.s^-1^.) during one hour and subsequently germinated in dark at 21°C for 96 hours, unless otherwise indicated. Abscisic acid was dissolved in ethanol and applied in final 2 µM concentration into the germination medium.

### In vitro cultures

To compare the phenotypes of *nramp2* CRISPR Cas9 mutants with the existing *nramp2-1* allele in Supplementary Figure 3, seeds were sown onto 0.6% (w/v) agar plates containing half-strength Murashige and Skoog medium supplemented with 1 % (w/v) sucrose and with or without addition of 20 µM MnSO_4_. The pH was adjusted to 5.7 with 2.5 mM MES-KOH. The seeds were stratified at 4°C and dark for 3 days and transferred to growth chamber for growth in a 16-light/8-h dark cycle 21°C with 65% relative humidity and a light intensity of 150 μmol.m^-2^. s^-1^.

### Luminescence assay

The seeds of *pZAT12::LUC* lines were sown under low intensity light (≈20 μmol.m^-2^ .s^-1^) in 384 well qPCR plates containing 0.5% (w/v) agar (pH 6 buffered with 2.5 mM MES-KOH) supplemented with 250µM luciferin. The plates were sealed with parafilm to avoid drying out and transferred into home-made dark chamber equipped with EMCCD camera (Andor iXon 897) and kept at 21°C. Luminescence was collected every 60 minutes for 96 h using 20 s exposure time. Luminescence signals were then quantified with Fiji (*43*) with mean values from ROI corresponding to each well. Luminescence values (L) were normalized as L/L₀, (L₀ = initial time point). ROS bursts were defined as three consecutive time points showed Δ(L/L₀) ≥ 0.2. Peak areas were quantified over 96 h using GraphPad Prism 10.6.1 for germinated seeds only (baseline = 1.5), excluding peaks < 0.5 luminescence units high or <3 adjacent points. Non-germinated seeds (showing < 2-fold L increase) were excluded, residual low signal seeds were verified by stereomicroscope at 96 h.

Curves were fitted using the Gompertz growth model: 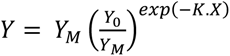

### Seed dissection

Seed dissection was performed according to Iwasaki and Molina (*44*). Briefly, dry seeds were imbibed in Milli-Q water for one hour, and embryo was released by applying gentle pressure on the seeds between two microscope slides. The mixture of embryo and the seed coat+endosperm was transferred into centrifuge tubes, 40% sucrose was added, mixed thoroughly, and centrifuged at 14.000 x g for 5 minutes. The physically separated embryos from the rest were rinsed three times with Milli-Q water and dried at 60°C in oven.

### Elemental analysis

Dry whole seeds and dissected seed parts were digested in solution of concentrated HNO_3_ and H_2_O_2_ (3:1 ratio) overnight at room temperature and consequently at 85°C for 24 hours. The samples were subsequently diluted with Milli-Q water. Elemental analysis was performed using inductively coupled plasma optical emission spectroscopy (ICP-OES; Agilent) using calibration standard solutions provided by the manufacturer. The ICP-OES instrument is calibrated for wavelength each time it is powered on, using a 5 ppm calibration solution provided by the manufacturer (ICP-OES & MP-AES Wave Cal 6610030100, Agilent Technologies). Performance tests (optics, sensitivity, and resolution) are then conducted. Quality control uses certified reference material (strawberry leaf powder, LGC7162), which closely matches the typical matrices analyzed. Emission wavelengths are selected based on precision from CRM measurements (±10% of theoretical values). The following elements were quantified at indicated wavelengths: calcium (Ca 393.366 nm), iron (Fe 238.204 nm), potassium (K 766.491 nm), magnesium (Mg 280.270 nm), manganese (Mn 257.610 nm), phosphorus (P 177.434 nm), sulfur (S 180.669 nm), and zinc (Zn 202.548 nm).

### Synchrotron X-ray fluorescence imaging

Whole intact seeds were used to perform sparse X-ray fluorescence (XRF) micro-tomography at NANOSCOPIUM beamline of SOLEIL Synchrotron, Saint Aubin. The process is described in Guo et al. (*45*). Briefly, 3D mesoscale XRF tomograms were collected at 11 keV using two energy-dispersive silicon drift detectors. Twenty projections were collected with a 18° angle over 360°. Each projection was collected with a 2 µm x 2 µm resolution and a 20 ms counting time. 3D volumes were reconstructed using the maximum-likelihood expectation–maximization (MLEM) algorithm. Then, high-resolution single slice tomography was collected with a 2 µm resolution, a 20 ms counting time and a 1° rotation angle, and reconstructed using the MLEM algorithm.

2D XRF imaging of dry seed sections was performed at LUCIA beamline of Synchrotron Soleil. Prior to imaging, dry seeds of *Arabidopsis thaliana* were embedded in optimal cutting temperature (OCT) medium, flash frozen in liquid nitrogen and cut into 25 µm sections using a cryo-microtome. The XRF was collected at 6.8 keV (Fig. 2g, Supplementary Fig. 5) or 7.2 keV (Supplementary Fig. 6) energy using 2.5 µm step size of at least four seed sections.

2D XRF imaging of fresh developing seeds was conducted at ID21 beamline of Synchrotron ESRF. Flowers were marked at anthesis and seeds of four siliques of three independent plants were dissected after 9 and 11 days after pollination, at bent cotyledon and mature green embryo stages, respectively. The seeds were transferred into OCT medium-containing PCR tube, frozen in liquid nitrogen and cut into 15 µm sections using a cryomicrotome. XRF was collected at 9.8 keV with 2.5 µm, 300 nm and 400 nm step sizes of seed sections originating from at least five individual seeds.

Elemental maps were generated by fitting the XRF spectra of each pixel through peak deconvolution using the PyMCA software and exported in 32-bit TIFF format. Mean pixel values for the embryo and seed coat regions of interest were measured in ImageJ after image segmentation, using threshold values determined separately for each seed tissue. The same thresholds were subsequently applied to all genotypes and developmental stages. Background pixel values were subtracted from the images.

### Laser ablation-coupled ICP-MS

Dried seeds were ablated with a laser ablation system (LA NewWave UP-213, Fremont, CA, USA) coupled to an ICP-MS system (ICP-MS 7900, Agilent, Hachioji, Japan), as previously reported in Wojcieszek et al. (*46*).

Analysis of Mn was conducted using the following laser ablation settings: laser spot diameter, 10 µm; laser energy 25%; fluence 1.0 J cm^-2^; repetition rate, 20 Hz; and scan speed, 10 µm sec^-1^. The ablated matter was transported into the ICP with helium gas (500 mL min^-1^) and mixed in a T-connector with aerosol obtained using a Micromist nebulizer. The double-pass Scott spray chamber was removed prior to ICP-MS.

Manganese values were normalised by determining the ratio of the intensity in count per second between manganese (isotope Mn^55^) and carbon (isotope C^13^) as reference element.

### Histochemical staining

#### GUS staining – GUS activity, GUS reflection

Siliques were cut open on one side, vacuum-infiltrated for 2 min, and incubated overnight at 37°C in GUS staining solution (50 mM sodium phosphate buffer pH 7.0, 0.5 mM ferro/ferricyanide, 0.05% Triton X-100, 1 mM X-Gluc). They were then washed in 50 mM sodium phosphate buffer, destained in 0.2 M NaOH + 1% SDS (3 h) to remove tannins, rinsed 3× in Milli-Q water (5 min each), decolorized in 1.25% sodium hypochlorite (30 min), rinsed again 3× in Milli-Q water (5 min each), and cleared/stored in chloral hydrate:glycerol:water (4:1:2, w/v/v).

### Scanning electron microscopy

Seeds were observed with environmental scanning electron microscope at UMR IATE, Montpellier.

### Statistics and reproducibility

Statistically significant differences among multiple groups were analyzed using one-way ANOVA followed by Tukey’s post hoc test (*P* < 0.05) or, when appropriate, using a non-parametric approach with a Kruskal–Wallis test and post hoc comparisons (*P* < 0.05). Differences between two groups were assessed using an unpaired Student’s t-test. The plots and statistical analyses were performed using GraphPad Prism version 10. All experiments were repeated at least twice with similar outcomes, and representative data from one experiment are shown.

## Acknowledgements

We acknowledge the MRI imaging facility, member of the France-BioImaging national infrastructure supported by the French National Research Agency (ANR-10-INBS-04, “Investments for the future”), and PHIV-La Gaillarde facility and the SAME platform. Luminescence signals were recorded with a custom BIOCORP setup (CNRS MITI interdisciplinary funding). Furthermore, SEM facility at UMR IATE, particularly the help of Aurélie Putois for the SEM imaging. We thank the Agence Nationale de la Recherche (Project ‘DEFIMAN’, ANR-19-CE20-0009-02). We acknowledge SOLEIL for provision of synchrotron radiation facilities beamlines LUCIA and NANOSCOPIUM and Synchrotron ESRF, beamline ID21.

## Author information

Authors and Affiliations

IPSiM, Univ Montpellier, CNRS, INRAE, Institut Agro, Montpellier, France

Alexandra Leskova, Tou Cheu Xiong, Sandrine Chay, Stéphane Mari, Catherine Curie

Institute of Interdisciplinary Research on Environment and Materials (IPREM), Université de Pau et des Pays de l’Adour, 64000 Pau, France

Marie-Pierre Isaure, Katarzyna Bierla

Synchrotron SOLEIL, L’Orme des Merisiers, 91190 Saint-Aubin, France UAR 1008 TRANSFORM, INRAE, 44316, Nantes, France

Camille Rivard

European Synchrotron Radiation Facility, Avenue des Martyrs, CS 40220, 38043 Grenoble, France

Hiram Castillo-Michel

Synchrotron SOLEIL, L’Orme des Merisiers, 91190 Saint-Aubin, France

Kadda Mejoubi, Andrea Somogyi

---

**Supplementary Fig. 1.**
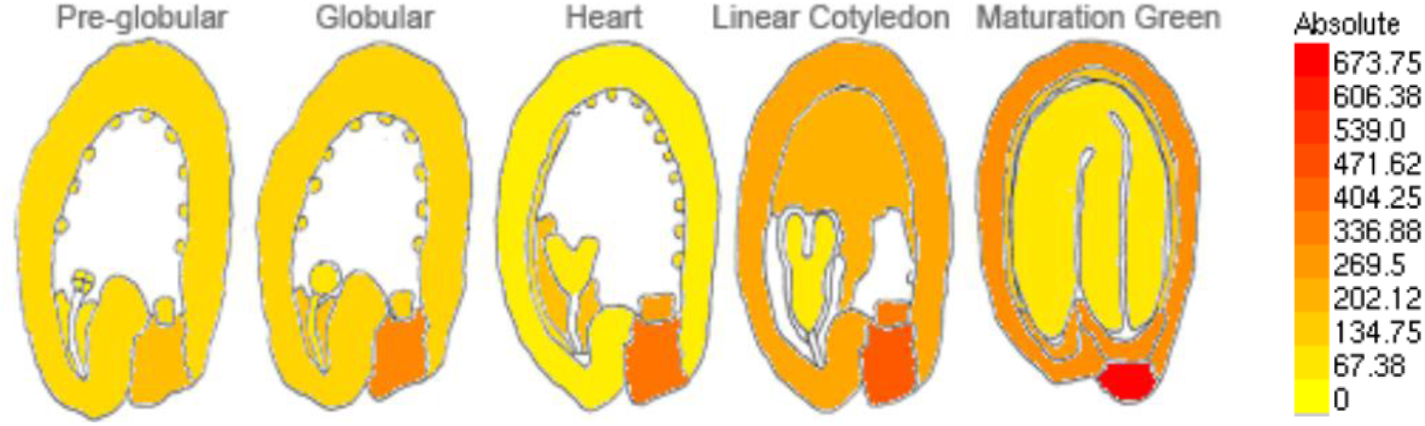
Tissue-specific enrichment of *NRAMP2* transcripts during seed development, according to Belmonte et al. (*19*).

**Supplementary Fig. 2.**
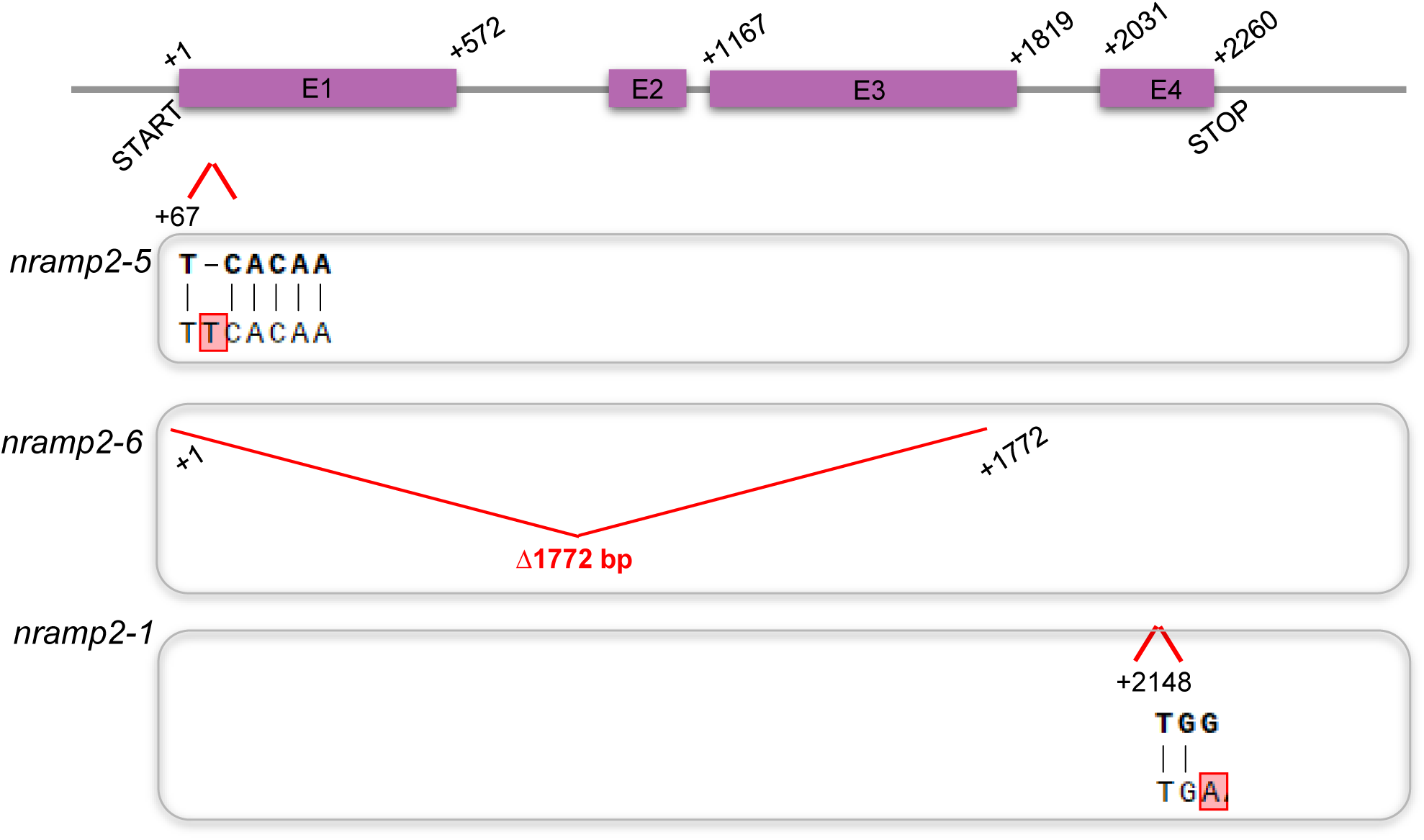
DNA sequences of the generated CRISPR CAS9 mutants. The *nramp2-5* allele carries a thymine insertion in the first exon, leading to frame shift and premature stop codon, and *nramp2-6* carries about a 1.8 Kb deletion of almost three entire exons. Base substitution in *nramp2-1* mutant described in Gao et al. (*18*).

**Supplementary Fig. 3.**
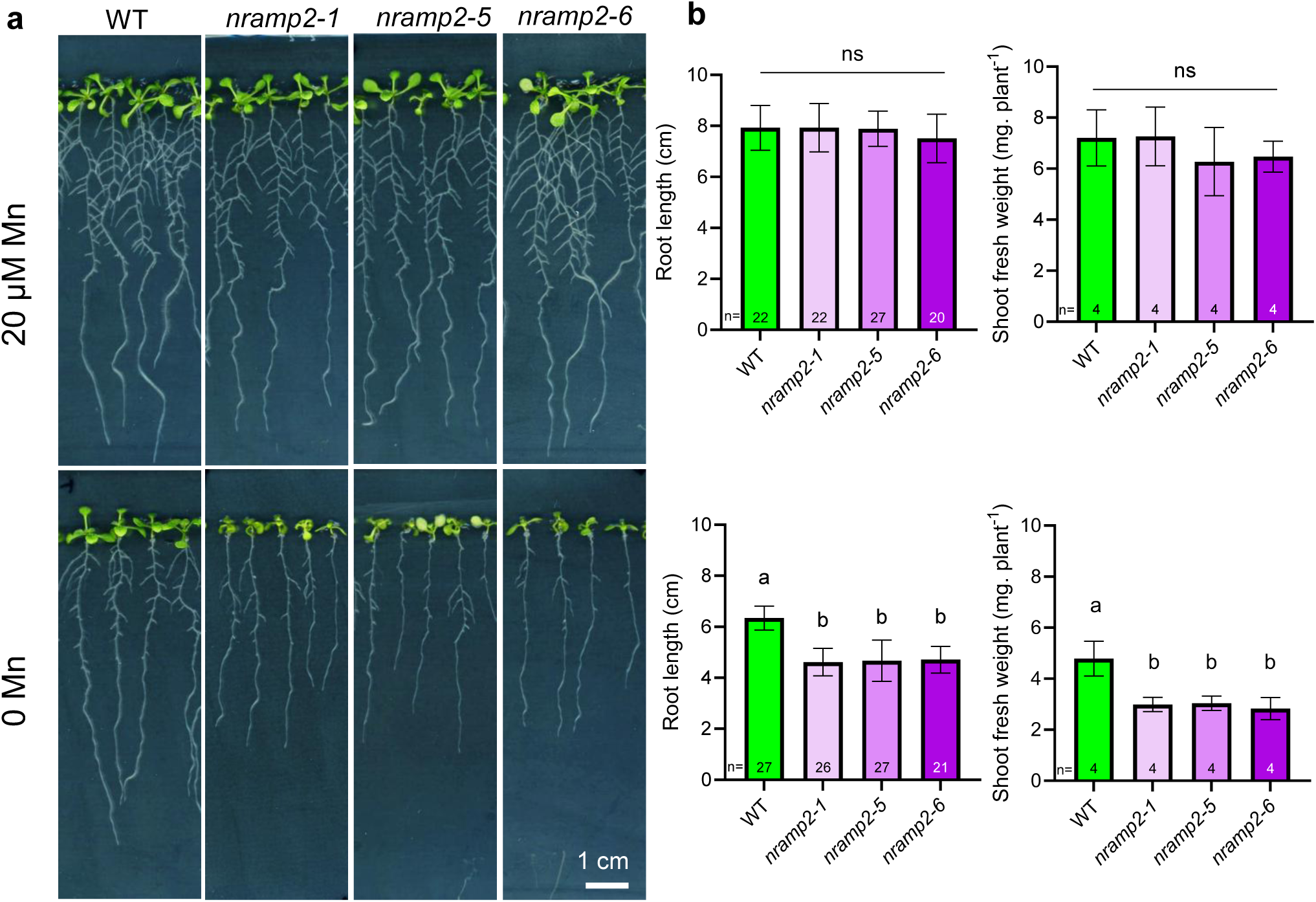
Phenotype of CRISPR CAS9 and already described *nramp2* mutants and its crosses. **a**, Visible phenotype of plantlets grown in ½ MS (20 µM Mn) or ½ MS with no added Mn for 10 days in long-days-condition. **b,** Root length and shoot biomass of plants grown in ½ MS with or without Mn. Bars represent means ± SD. Different letters indicate significant differences (one-way ANOVA followed by post-hoc Tukey’s test, *P* < 0.05).

**Supplementary Fig. 4.**
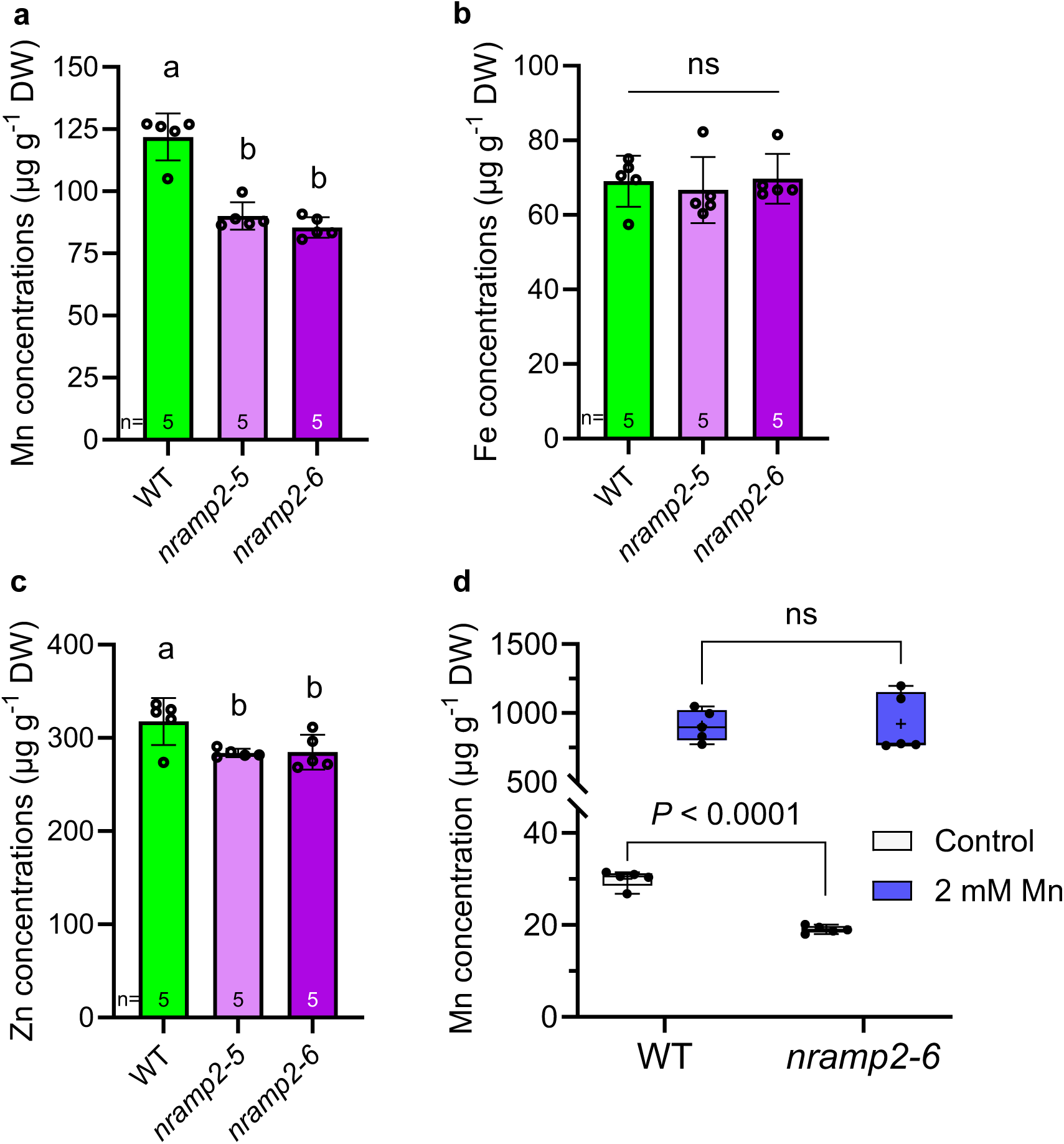
Mn accumulation is reduced in *nramp2*. Content of **a,** Mn, **b**, Fe and **c,** Zn in dry seeds analysed by ICP-OES. Bars represent means ± SD. Different letters indicate significant differences (one-way ANOVA followed by post-hoc Tukey’s test, *P* < 0.05). **d,** Contents of Mn in seeds originating from plants watered with or without 2 mM MnSO_4_ throughout their development. Elemental concentrations were measured from seeds harvested at the end of the plants’ life cycle. Plants were potted into standard soil substrate of the greenhouse, and watered weekly with tap water unless indicated otherwise. Box plot represents all data points, whiskers the 25th percentile, median and 75th percentile. *P*-values were determined by two-tailed unpaired Student’s *t* test.

**Supplementary Fig. 5.**
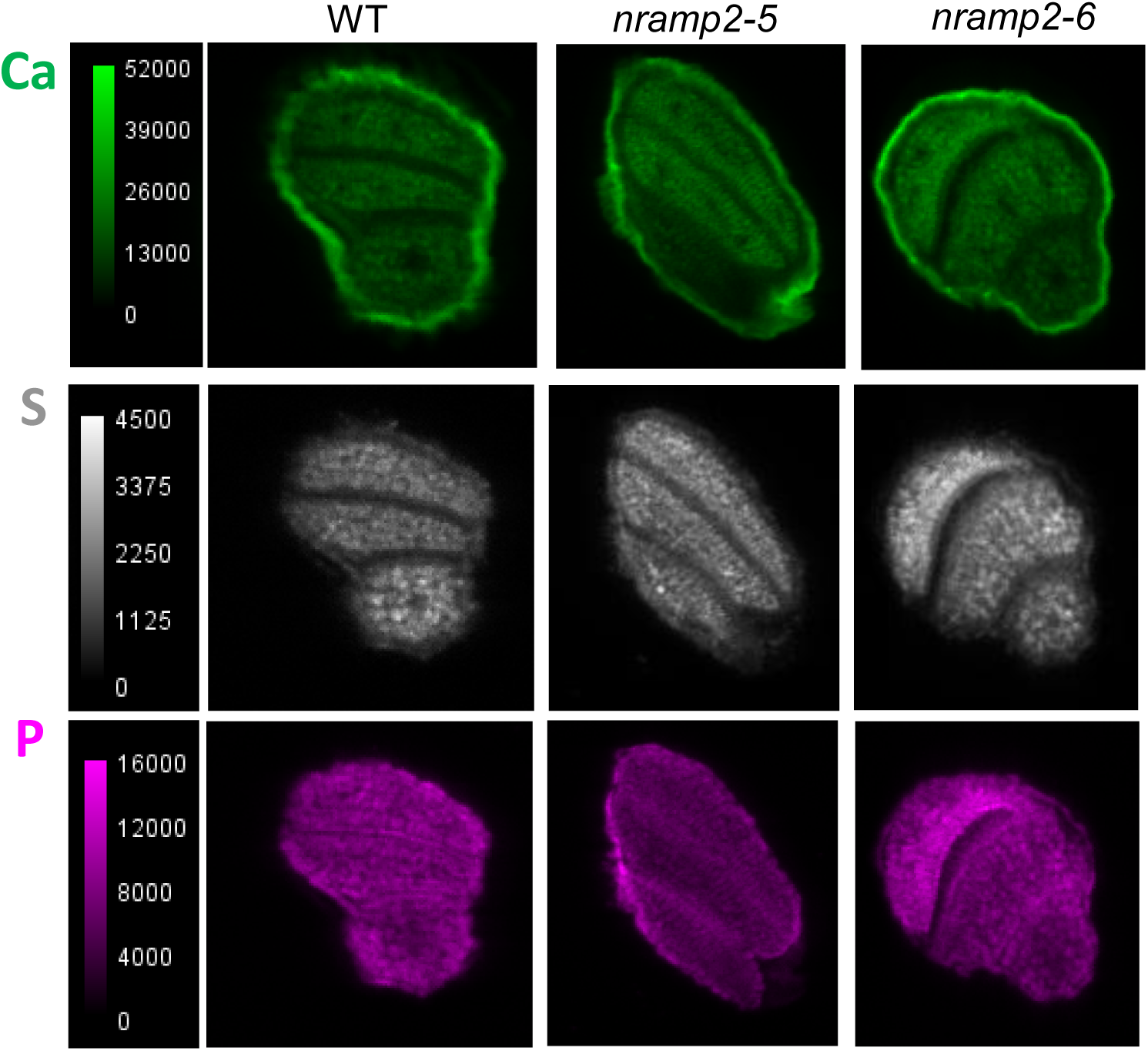
Macroelement distribution in dry seed sections illustrated in Figure 2. Micro X-ray fluorescence imaging of calcium (Ca), sulfur (S) and phoshorus (P) using Lucia beamline of Synchrotron Soleil. Representative images of seed sections are shown.

**Supplementary Fig. 6:**
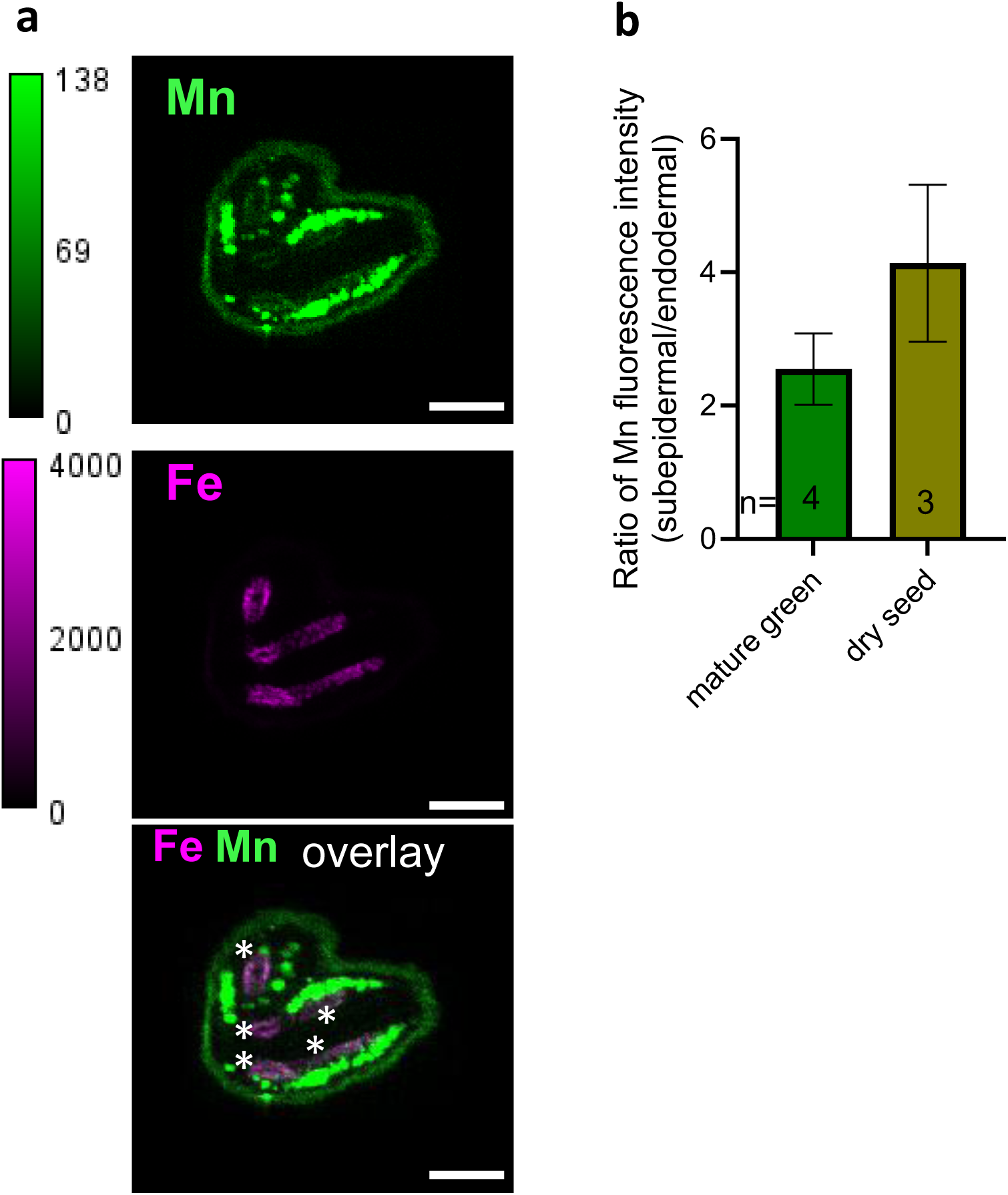
Manganese accumulation in dry seeds of wild type Arabidopsis plants. **a,** Endodermal Mn is still detectable in the dry seeds. Asterisks mark the endodermal hotspots, colocalizing with Fe. **b,** Proportion of Mn fluorescence accumulating in subepidermal vs endodermal vacuoles. Scalebars = 100 µm.

**Supplementary Fig. 7:**
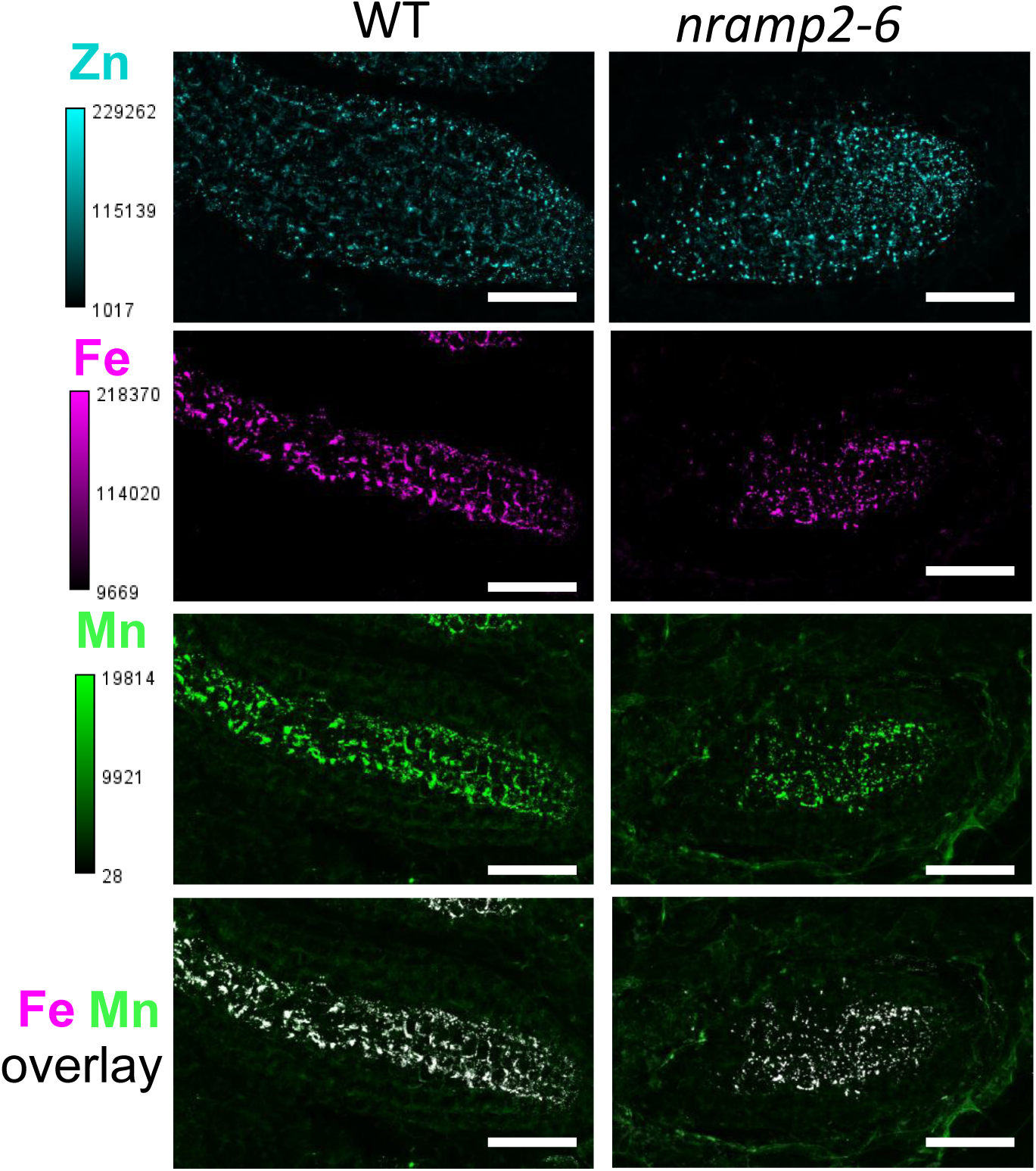
Metal accumulation patterns in bent cotyledon-staged seeds. The images are a crop of the embryo radical. Scalebars: 400 nm.

**Supplementary Fig. 8.**
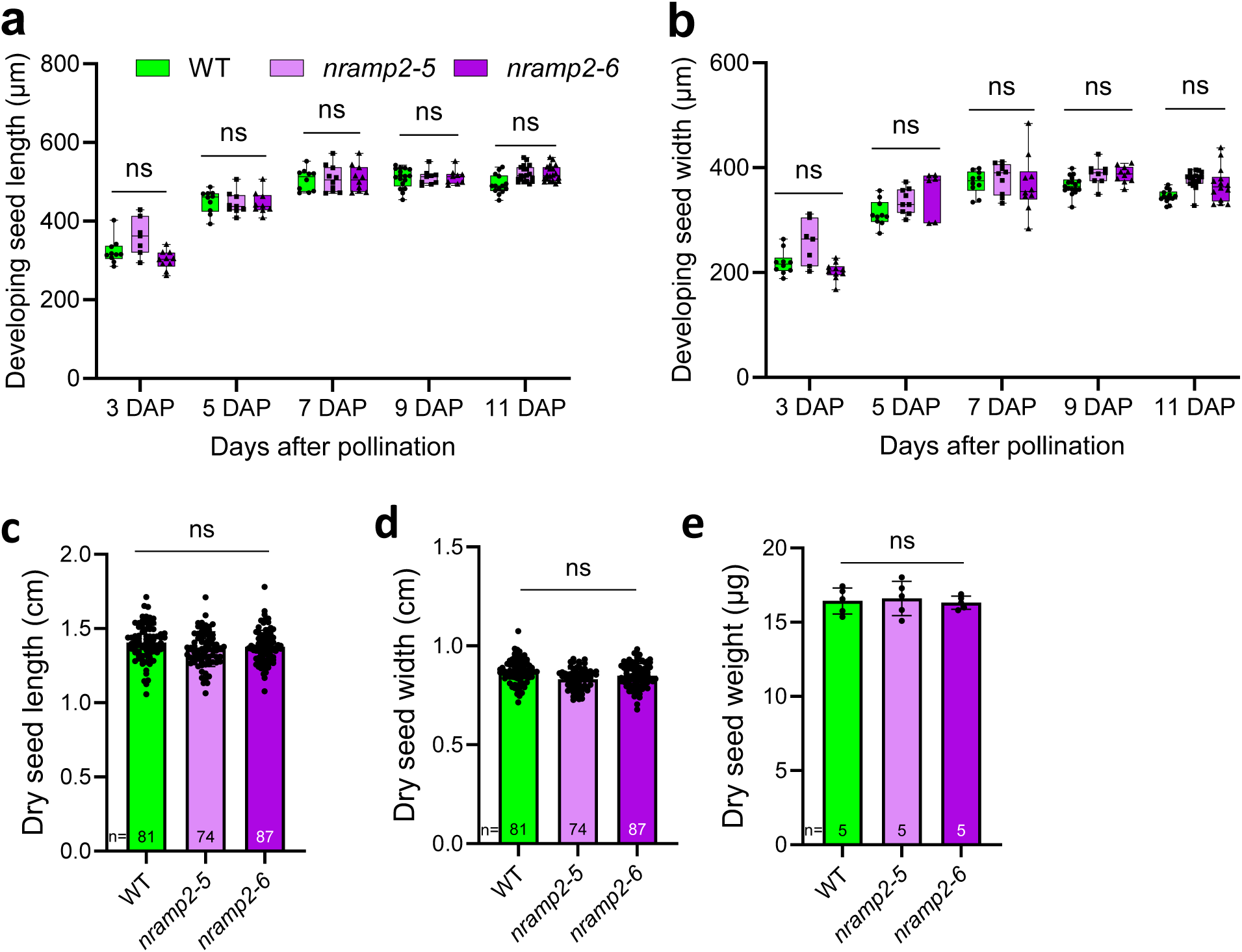
Loss of *NRAMP2* does not evoke changes in seed development and dry seed parameters. **a,b,** Length and width of developing seeds from 3 to 11 days after pollination (DAP). Box plot represents all data points, whiskers the 25th percentile, median and 75th percentile. Number of replicates ≈ 10. **c,d,e,** Length, width and weight of dry seeds. Statistical significance was determined by one-way ANNOVA followed by Tukey’s test.

**Supplementary Fig. 9.**
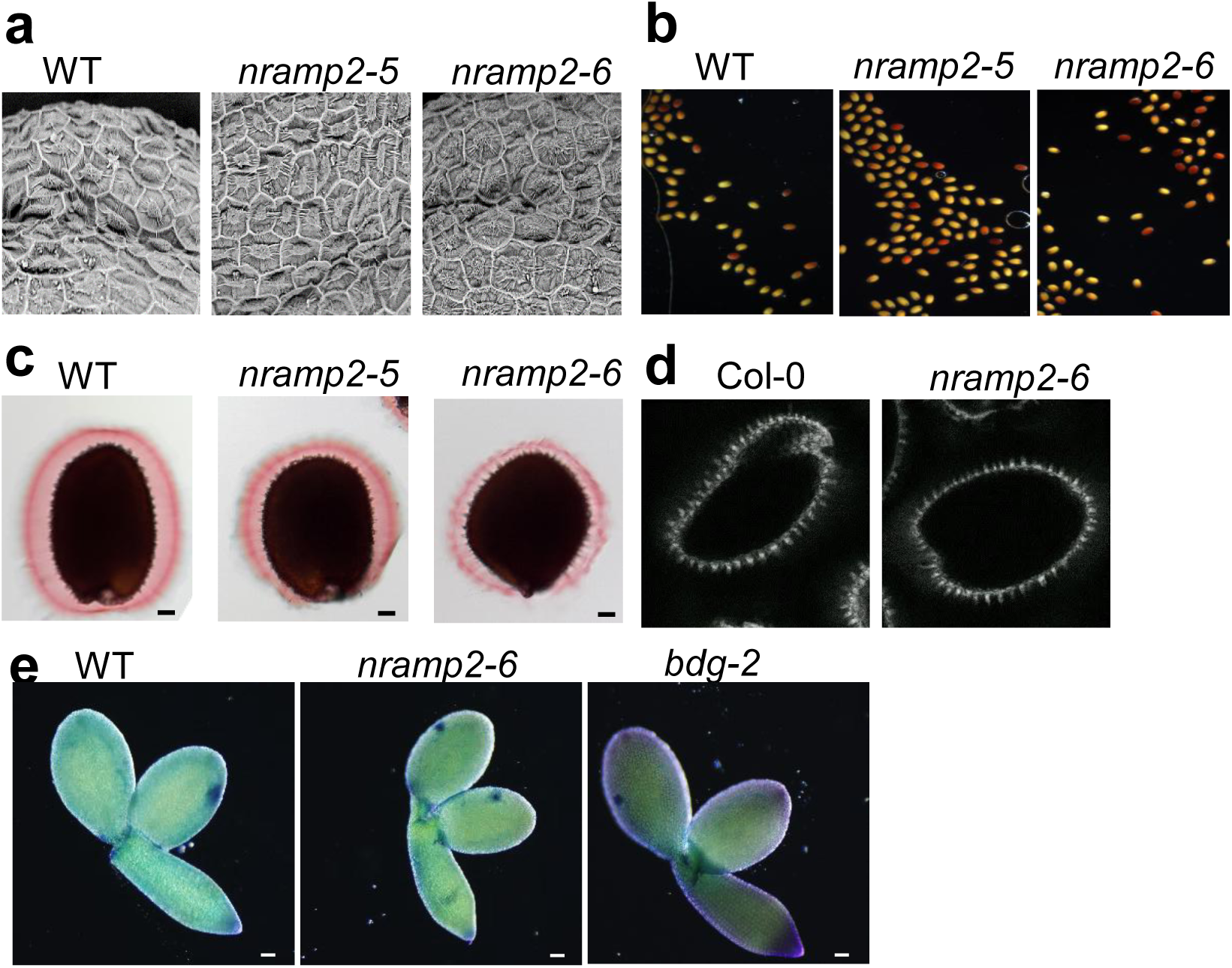
Mutation of *NRAMP2* does not have a major impact on seed coat structure and cell wall production. **a,** Scanning electron microscopy of seed coat of dry seeds. **b,** Seed coat permeability assay by tetrazolium staining of imbibed seeds, **c,** Ruthenium red staining of pectin in imbibed seeds, **d,** Calcofluor staining of cellulose of imbibed seeds, **e,** Toluidine blue staining of cuticle of dissected embryos (11DAP). Scalebars= 50 µm.

**Supplementary Fig. 10:**
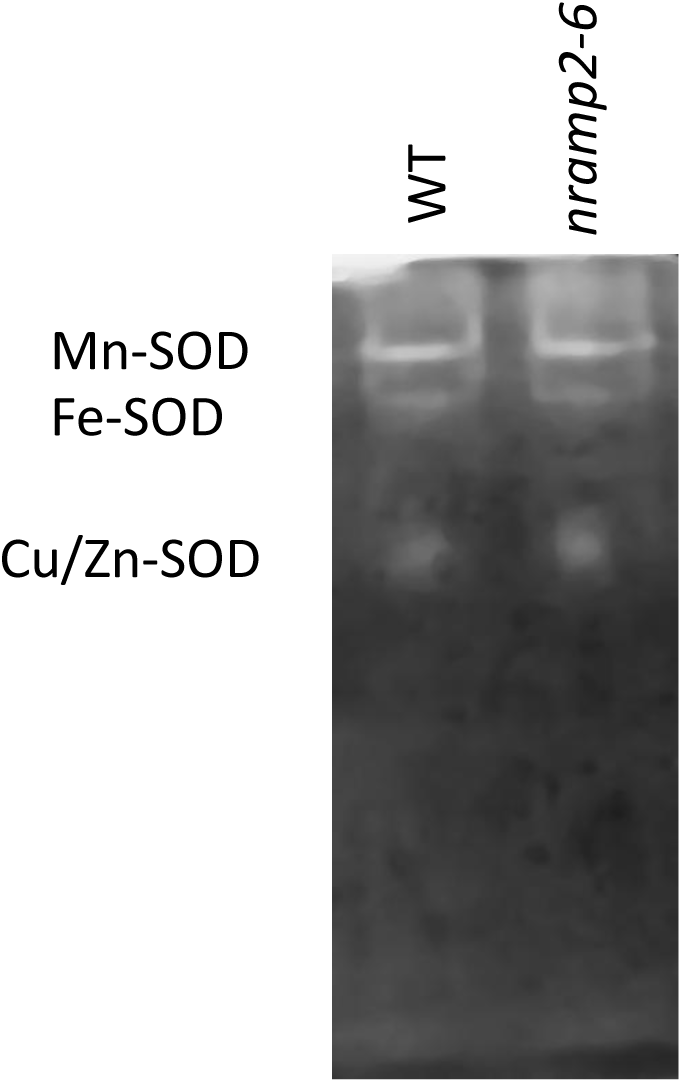
Mn-SOD activity is not altered in developing *nramp2* mature seeds.

**Supplementary Table I.** Elemental concentrations in dry seeds. Values represent the means ± SD of five independent samples. The letters indicate statistical significance among genotypes at *P* < 0.05 according to one-way ANNOVA and Tukey’s post-hoc test.

| Element | Elemental concentrations ($\mu\text{g.g DW}^{-1}$) | | | | | |
| --- | --- | --- | --- | --- | --- | --- |
|  | WT |  | *nramp2-5* |  | *nramp2-6* |  |
| Ca | 4602.2 | ± 299.1 a | 4584.3 | ± 97.4 a | 4750.6 | ± 257.9 a |
| K | 27083.3 | ± 3223.9 a | 27987.8 | ± 1409.4 a | 26770.2 | ± 3910.1 a |
| Mg | 2770.5 | ± 25.9 a | 2928.1 | ± 75.1 a | 2980.1 | ± 187.5 a |
| P | 9726.7 | ± 838.8 a | 10657.7 | ± 280.9 a | 10566.0 | ± 767.2 a |
| S | 13294.6 | ± 588.1 a | 12378.1 | ± 111.3 b | 13161.4 | ± 570.5 ab |

